## Supplementary info for "Repurposing host-guest chemistry to sequester virulence and eradicate biofilms in multidrug resistant *Pseudomonas aeruginosa* and *Acinetobacter baumannii*"

**Supplementary Table 1. Macrocycles used in the HSL screening.**

| Full name | Cavity Size (Å) | Molecular Weight (M) | Dissolved in / miscible with (M) |
|---|---|---|---|
| 12-Crown-4 | 1,50 | 172,00 | 1X PBS (M) |
| 2-hydroxymethyl-12-crown-4 | 2,20 | 206,00 | 1X PBS (M) |
| 1-Aza-15-Crown-5 | 2,20 | 219,00 | 1X PBS |
| 15-Crown-5 | 2,20 | 219,00 | 1X PBS (M) |
| 4-sulfocalix[4]arene | 3,00 | 744,00 | 1X PBS |
| Calix[4]resorcinarene | 3,00 | 1159,00 | 1X PBS |
| 18-Crown-6 | 3,20 | 264,00 | 1X PBS |
| Cucurbit[6]uril hydrate | 3,90 | 996,00 | 1X PBS |
| Pillar[5]arene | 4,60 | 2260,00 | 1X PBS |
| (2-hydroxypropyl)-α-cyclodextrin | 5,70 | 1180,00 | 1X PBS |
| α-cyclodextrin | 5,70 | 972,00 | 1X PBS |
| Calix[6]arene | 7,60 | 636,00 | DMSO |
| 4-tert-butylcalix[6]arene | 7,60 | 973,00 | DMSO |
| Methyl-β-cyclodextrin | 7,80 | 1320,00 | 1X PBS |
| (2-hydroxypropyl)-β-cyclodextrin | 7,80 | 1396,00 | 1X PBS |
| β-cyclodextrin | 7,80 | 1135,00 | 1X PBS |
| 2-hydroxypropyl-ɣ-cyclodextrin | 9,50 | 1762,00 | 1X PBS |
| ɣ-cyclodextrin | 9,50 | 1297,00 | 1X PBS |
| Calix[8]arene | 11,70 | 849,00 | DMSO |

**Supplementary Table 2. Plasmids used in this study.**

***Receptor plasmids***

| Identifier | Description | Source |
|---|---|---|
| PC21 *pJ23100-LuxR* | In-house | (CJ)[1] |
| PC22 *pJ23100-CinR* | In-house | (CJ)[2] |
| PC72 *pJ23100-RhlR* | In-house | (PS) *P. aeruginosa* strain ATCC15692 genome |
| PC125 *pJ23100-RpaR* | In-house | (PS) *R. palustris* strain CGA009 genome |

*Promoter plasmids*

| Identifier | Description | Source |
|---|---|---|
| PC71 *pLux-eGFP* | In-house | (PS)[1] |

| | | |
|---|---|---|
| PC86 *pRhl(NM)-B0034-eGFP* | In-house | (PS) BBa_K1949060 + RBS_BBa_B0034 |
| PC87 *pRhl(RR)-B0034-eGFP* | In-house | (PS) BBa_K1529320 + RBS BBa_B0034 |
| PC90 *pRpa-eGFP* | In-house | (PS) Synthetic[1,3] + pLux RBS[4] |
| PC136 *pRhl(RR)-eGFP* | In-house | (PS) BBa_K1529320 + pLux RBS[1] |
| PC138 *pCin-eGFP* | In-house | (PS) + pLux RBS[1] |

**Supplementary Table 3. Clinical isolates used in this study.**

Attached as separate excel.

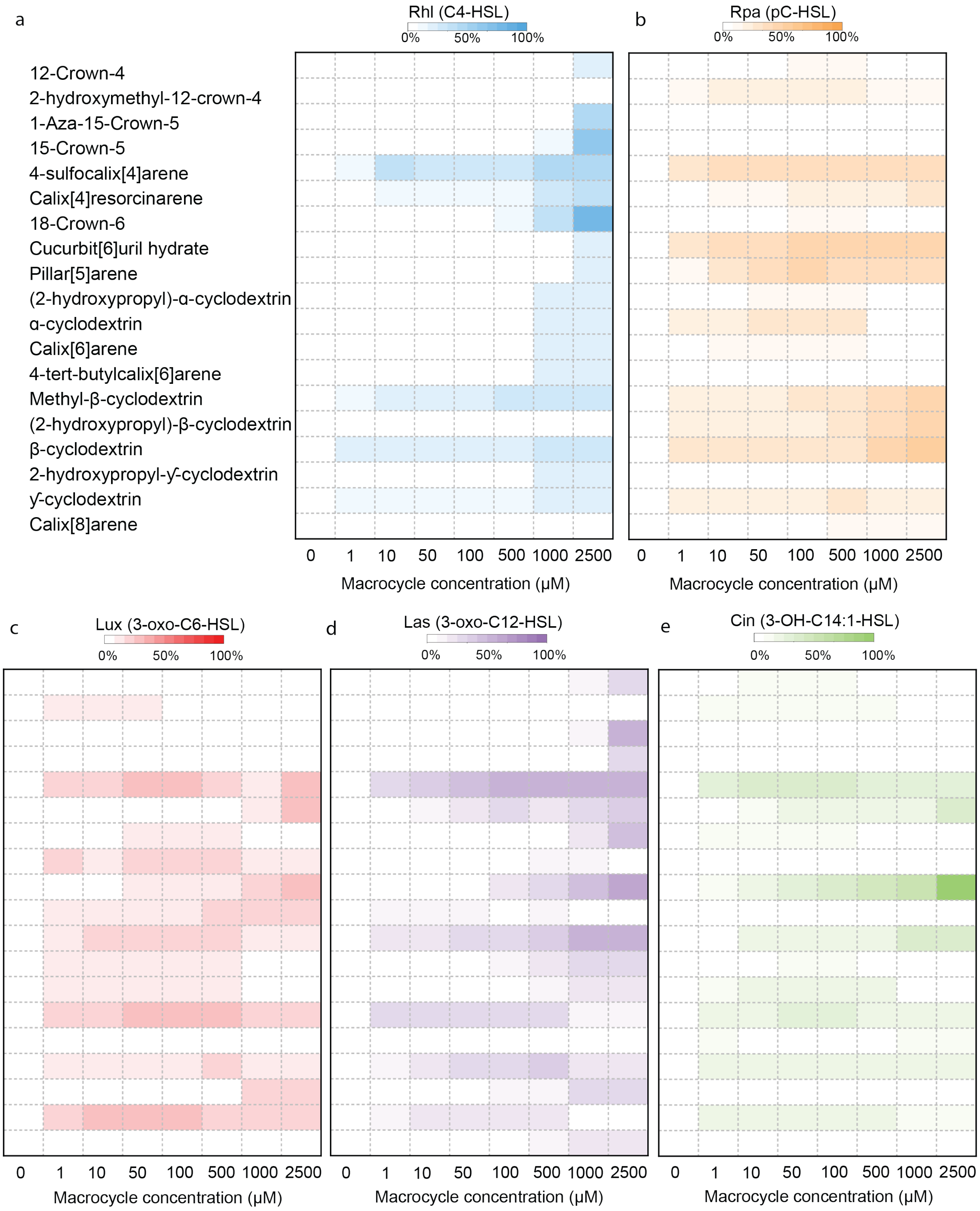

**Supplementary Figure 1.** Heatmaps depicting the interaction between a set of 19 different macrocycles selected and 5 HSLs, C4 (**a**), pC (**b**), 3-Oxo-C6 (**c**), 3-Oxo-C12 (**d**) and 3-OH-C14:1 HSL (**e**). Macrocycle coding correlates to Supplementary Information Table 1.

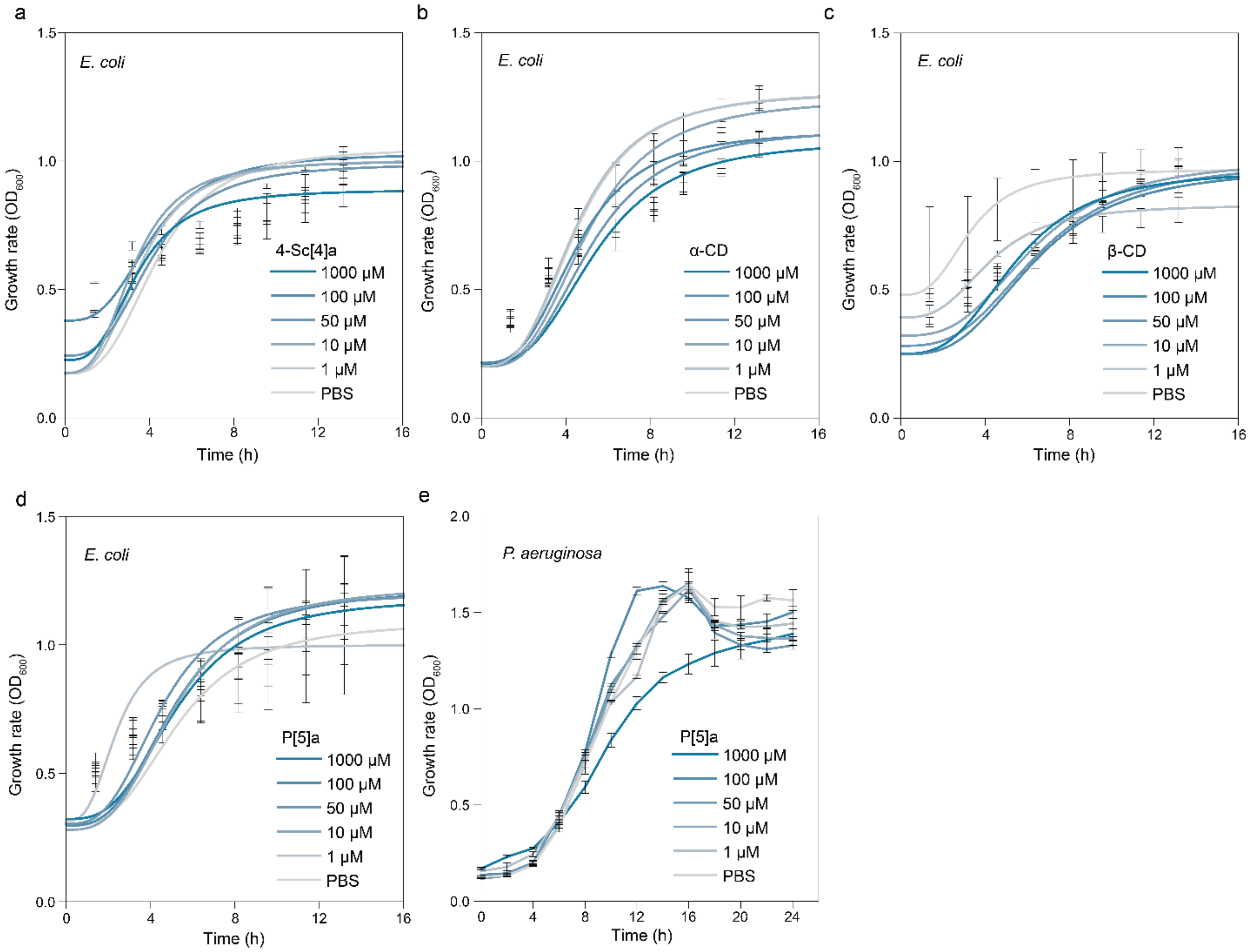

**Supplementary Figure 2.** Effect of selected macrocyclic hosts on bacterial growth rates. **a**) Effects of increasing concentrations of 4-Sc[4]a, **b**) α-CD, **c**) β-CD and **d**) P[5]a on the growth rate ($OD_{600}$) of the *E.coli* biosensor screen. The data represent average values of biological replicates ± s.d (n=3 per group). **e**) Effects of increasing concentrations of P[5]a on the growth rate ($OD_{600}$) of the *P. aeruginosa*. An increase in $OD_{600}$ is observed in lower concentrations of P[5]a (1-100 μM) and the PBS control, which is due to the production of the toxin pyocyanin, affecting the $OD_{600}$ measurement. In the 1000 μM P[5]a concentration, the toxin production is suppresses. As such, the 1000 μM concentration shows a standard growth curve. The data represent average values of biological replicates ± s.d (n=3 per group).

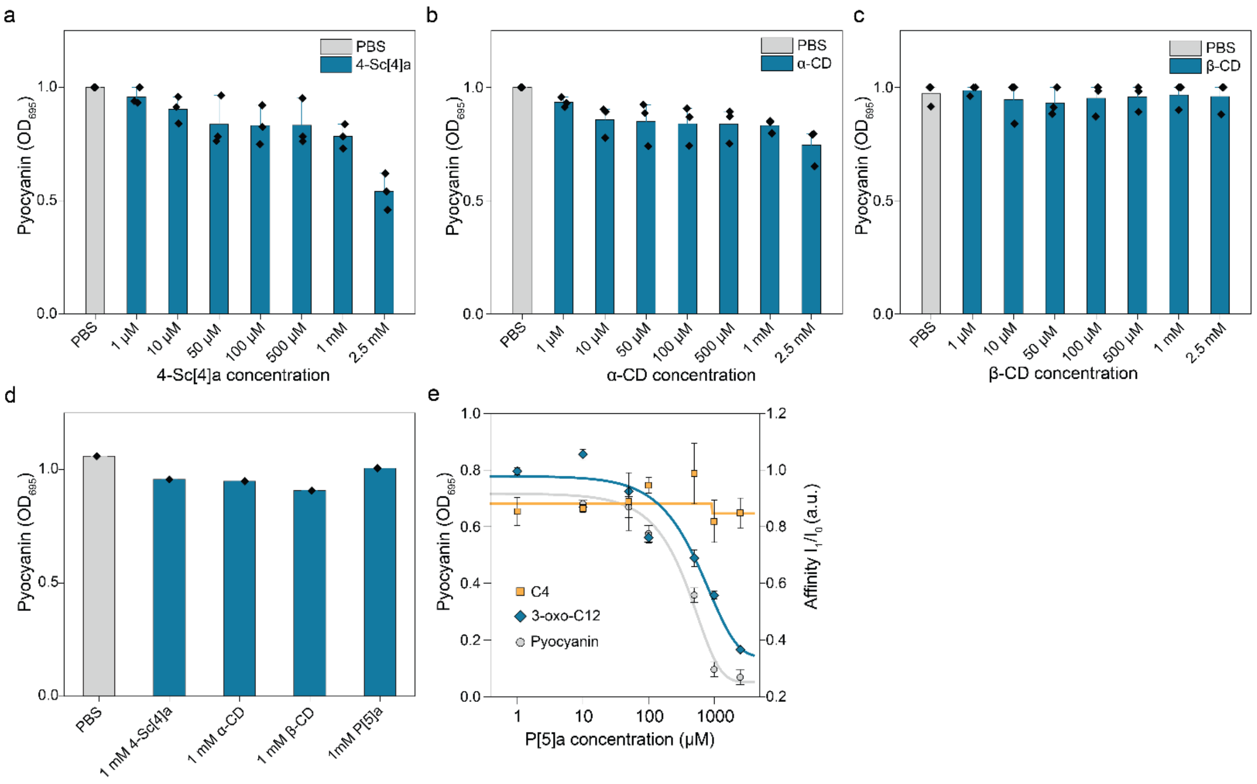

**Supplementary Figure 3.** Effect of selected macrocyclic hosts on pyocyanin production in *Pseudomonas aeruginosa* PAO1. **a**) Effects of increasing concentrations of 4-Sc[4]a, **b**) α-CD and **c**) β-CD on pyocyanin levels after a 24h incubation. The data represent average values of biological replicates ± s.d (n=3 per group). **d**) Effect of selected macrocyclic hosts on purified culture fluid of untreated PAO1 with high levels of pyocyanin, incubated for 24h shows that the macrocyclic hosts do not directly interact with the toxin pyocyanin. The data represent average values of biological replicates ± s.d (n=3 per group). **e**) Comparison between the dose-dependent effects of P[5]a on the production of pyocyanin (grey line) and the dose-dependent effects of P[5]a on the C4 (orange) and 3-oxo-C12 (blue) HSLs (the HSL signaling systems of *P. aeruginosa)* in the fluorescent *E. coli* reporter screen The data represent average values of biological replicates ± s.d (n=3 per group).

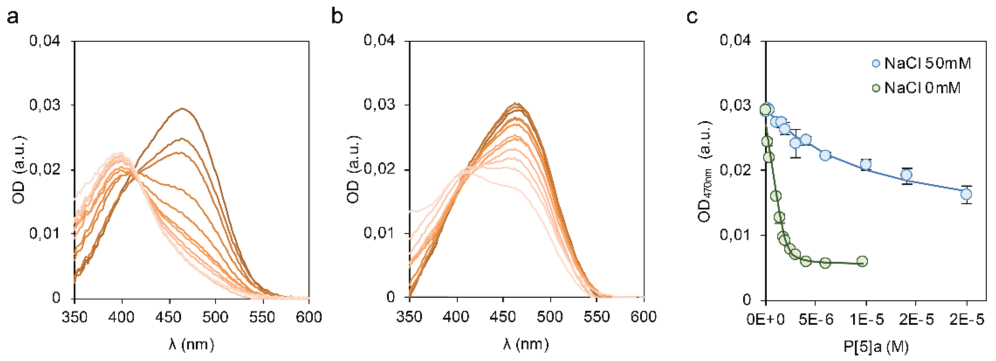

**Supplementary Figure 4.** Absorption spectra of MO ($2.0x10^{-6}$ M) titrated with increasing amounts of P[5]a in **a**) 0 and **b**) 50 mM of NaCl. **c**) Absorption at 470 nm at 0 and 50 mM of NaCl, and the fitted curves for a 1:1 host-guest model. The data represent average values of biological replicates ± s.d (n=3 per group).

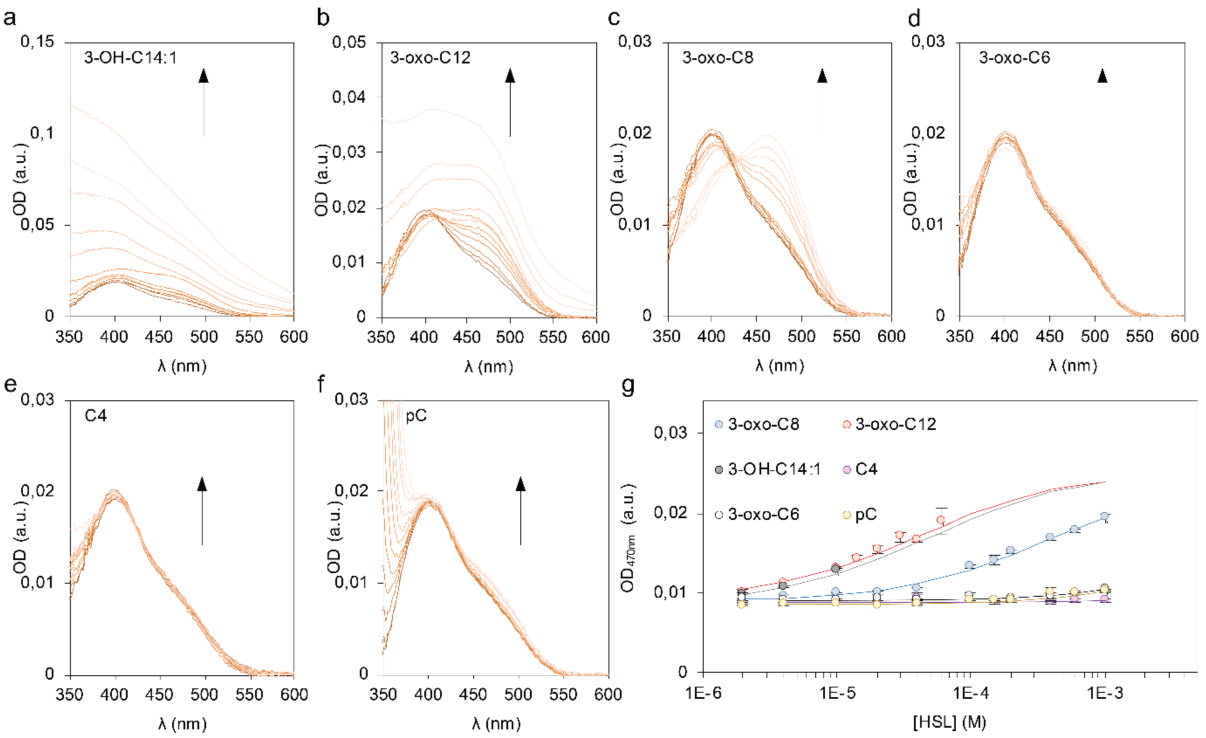

**Supplementary Figure 5**. Absorption spectra of a MO-P5a solutions ([MO] = [P[5]a] = $2.0x10^{-6}$ M; [NaCl] = 0 mM; DMSO = 1%), titrated with increasing amounts of HSLs. **a**) [3-OH-C14:1 HSL]: 0 → $2.0x10^{-4}$ M, **b**) [3-oxo-C12 HSL]: 0 → $2.0x10^{-4}$ M, **c**) [3-oxo-C8 HSL]: 0 → $1.0x10^{-3}$ M, **d**) [3-oxo-C6 HSL]: 0 → $1.0x10^{-3}$ M, **e**) [C4 HSL]: 0 → $1.0x10^{-3}$ M, **f**) [pC HSL]: 0 → $1.0x10^{-3}$ M. **g**) Absorption at 470 nm of the solutions, plotted as average of the triplicates. Solid line of corresponding color shows the fitted model. The data represent average values of biological replicates ± s.d (n=3 per group).

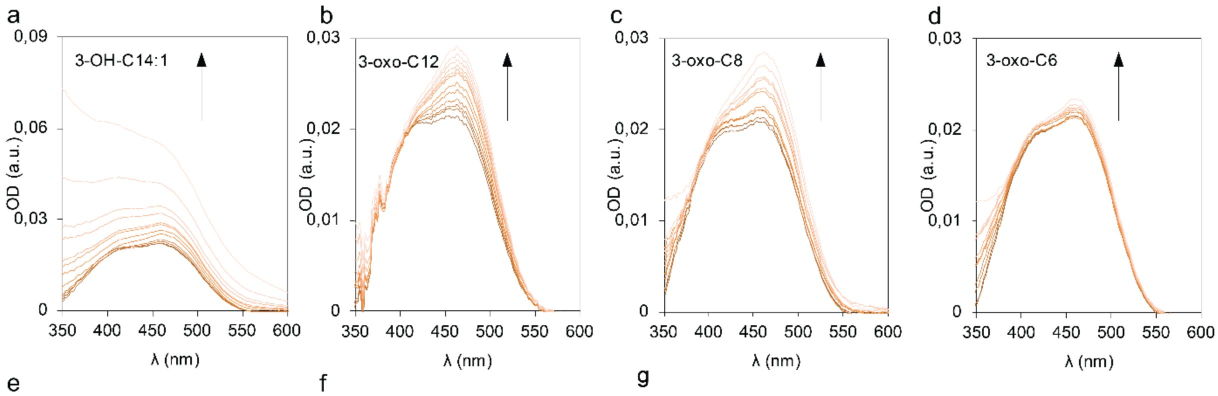

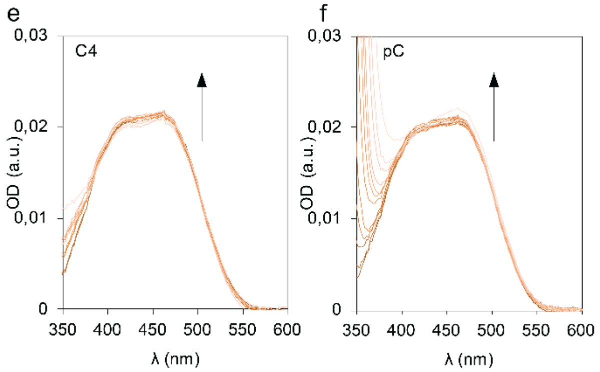

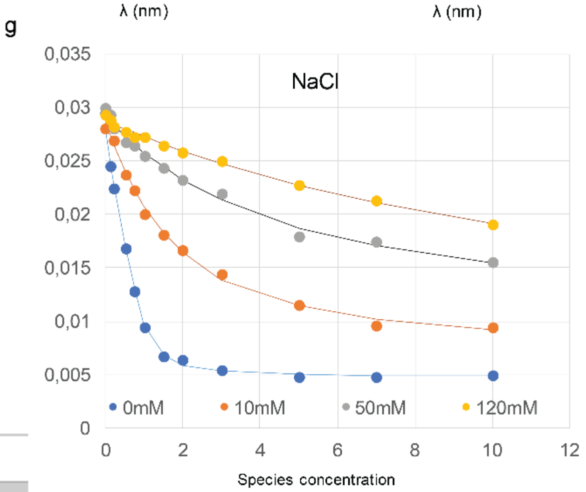

h

| [NaCl] (mM) | K2 ($M^{-1}$) |
|---|---|
| 0 | $(8.30\pm0.07)x10^6$ |
| 10 | $(4.53\pm0.03)x10^5$ |
| 50 | $(1.37\pm0.05)x10^5$ |
| 120 | $(3.94\pm0.05)x10^4$ |

**Supplementary Figure 6.** Absorption spectra of a MO-P[5]a solutions ([MO] = $2.0x10^{-6}$ M; [P[5]a] = $1.0x10^{-5}$ M; [NaCl] = 50 mM; DMSO = 1%), titrated with increasing amounts of HSLs. **a**) [3-OH-C14:1 HSL]: 0 →$1.0x10^{-4}$ M, **b**) [3-oxo-C12 HSL]: 0 →$1.0x10^{-4}$ M, **c**) [3-oxo-C8 HSL]: 0 →$1.0x10^{-3}$ M, **d**) [3-oxo-C6 HSL]: 0 →$1.0x10^{-3}$ M, **e**) [C4 HSL]: 0 →$1.0x10^{-3}$ M, **f**) [pC HSL]: 0 →$1.0x10^{-3}$ M. **g**) Effect of increasing concentrations on the MO-P[5]a binding. **h**) Binding affinities for host interactions of MO-P[5]a in differing concentration of NaCl, as seen in (**g**). The data represent average values of biological replicates ± s.d (n=3 per group).

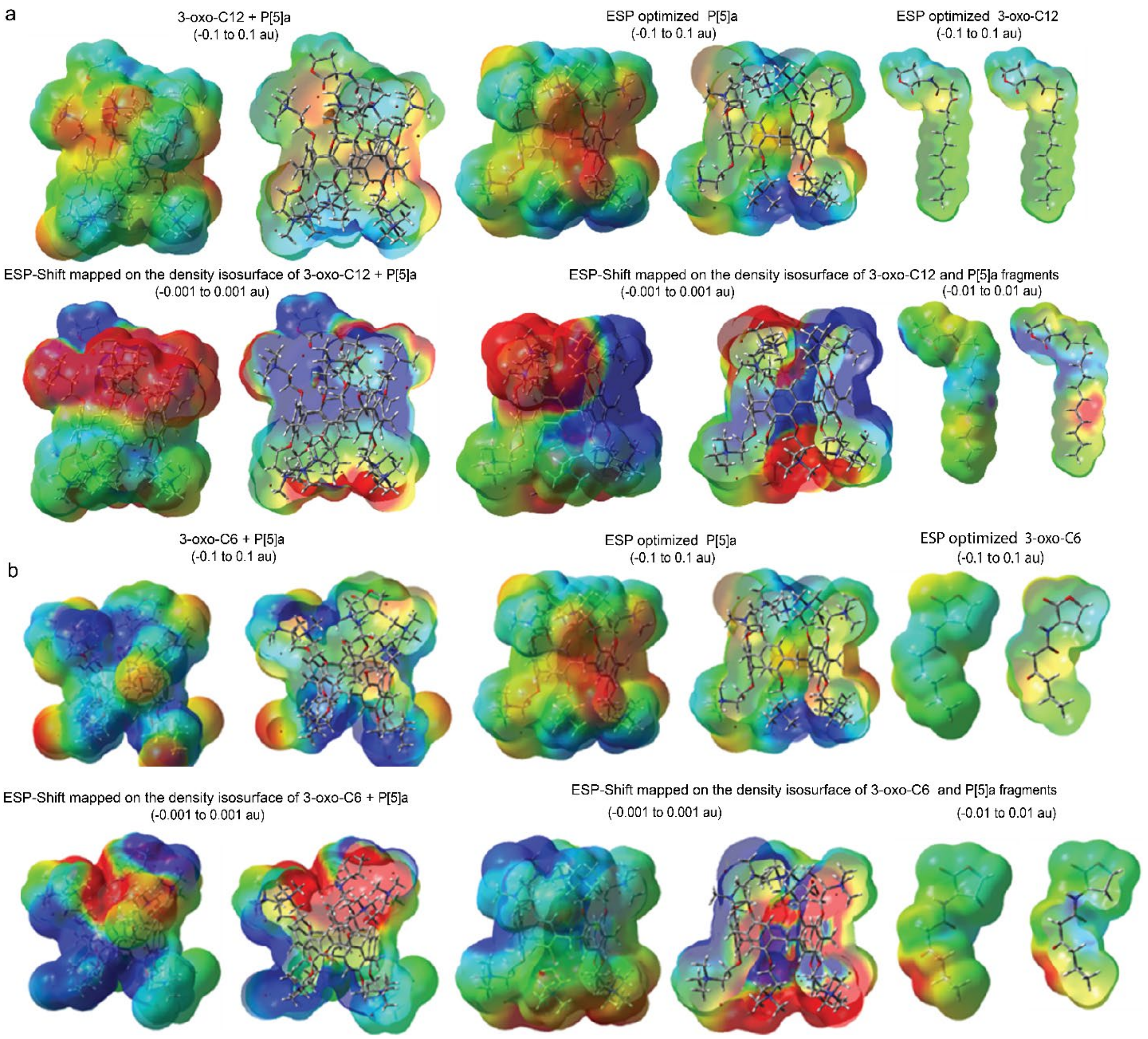

**Supplementary Figure 7.** Electrostatic potential and the electrostatic potential shift mapped on the molecular electron density surface of the optimized complex of **a**) 3-oxo-C12 and P[5]a, and the optimized structures of both the P[5]a and 3-oxo-C12 fragments in the complex; **b**) 3-oxo-C6 and P[5]a, and related surface of the optimized strucures of P[5]a and 3-oxo-C6 fragments. Note that deep blue is the most negative, deep red the most positive electron density. Note too the difference in scale between the absolute and shift figures.

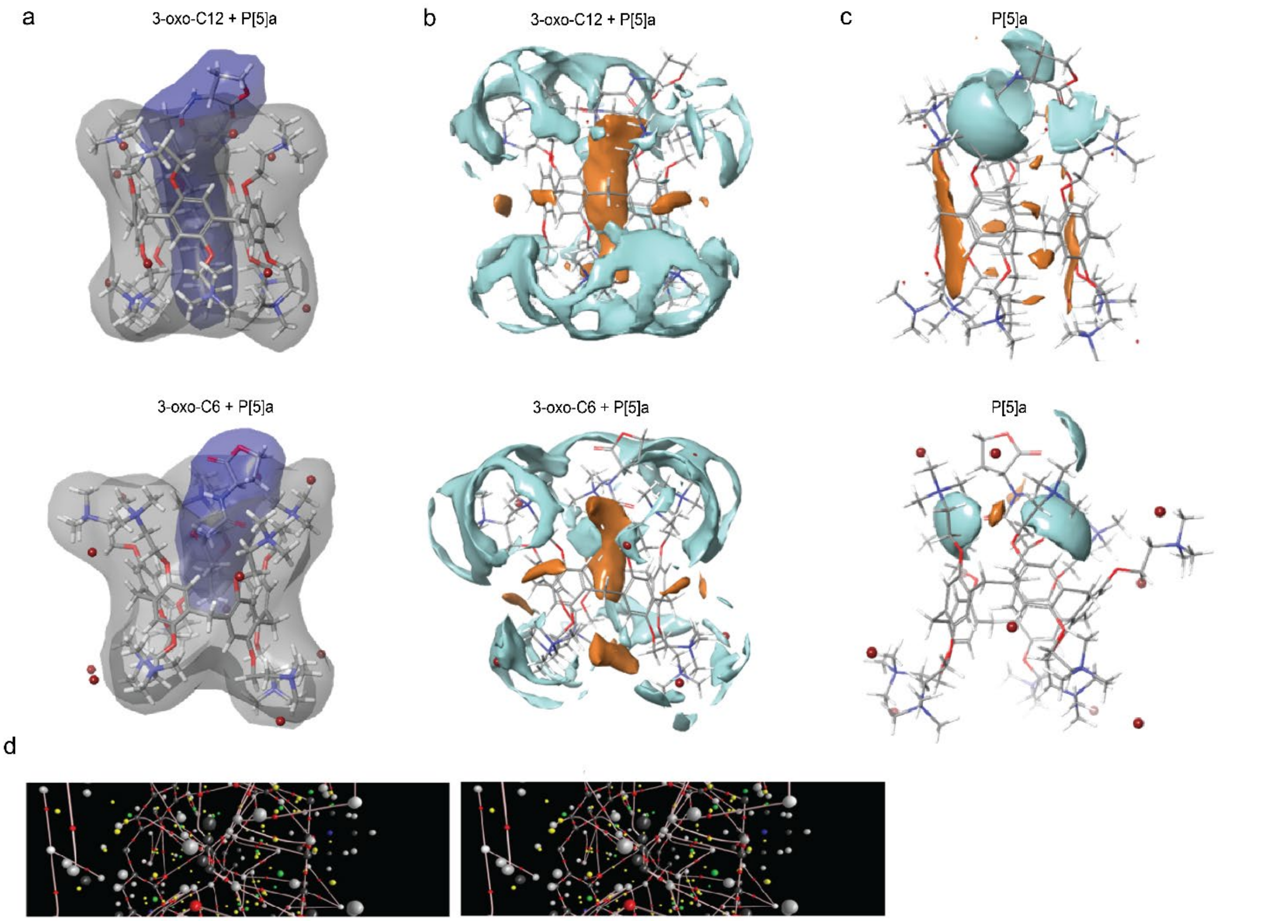

e

| | $E_{dist/P[5]}$ (kcal/mol) | $E_{int}$ (kcal/mol) | Binding Site Surface (Å$^2$) | Ligand Surface (Å$^2$) | $\sum\rho_{total\ au}$ | P[5] philic Surface (Å$^2$) | P[5] phobic Surface (Å$^2$) | Ligand philic Surface (Å$^2$) | Ligand phobic Surface (Å$^2$) |
|---|---|---|---|---|---|---|---|---|---|
| 3-oxo-C12 + P[5]a | 63.0 | -41.1 | 1184.83 | 297.04 | 0.08 | 731.910 | 124.898 | 136.100 | 70.459 |
| 3-oxo-C6 + P[5]a | 71.0 | -28.5 | 1201.21 | 197.61 | 0.04 ≥ | 494.741 | 117.650 | 109.716 | 6.255 |

**Supplementary Figure 8. a**) The calculated density isosurfaces of 3-oxo-C12, and 3-oxo-C6 bound to P[5]a; **b**) The calculated hydrophobic (orange) and hydrophilic (blue) surface interactions of the host in the presence of the guest, and **c**) guest in the presence of the host; both analyses were conducted using the Schrödinger suite of programs. **d**) The QTAIM molecular graph showing bond critical points and bond paths were obtained from the M062X/6-311G**electron density for the favored and 3-oxo-C12 complexes calculated using AIM2000; **e**) The estimated distortion and interaction energies, the binding site areas, the hydrophobic surface areas and the calculated total electron density at the bond critical points of the key vWd interactions in the binding cavity for the M062X/6-311G**electron density for the P[5]a and 3-oxo-C12, and P[5]a and 3-oxo-C6 complexes.

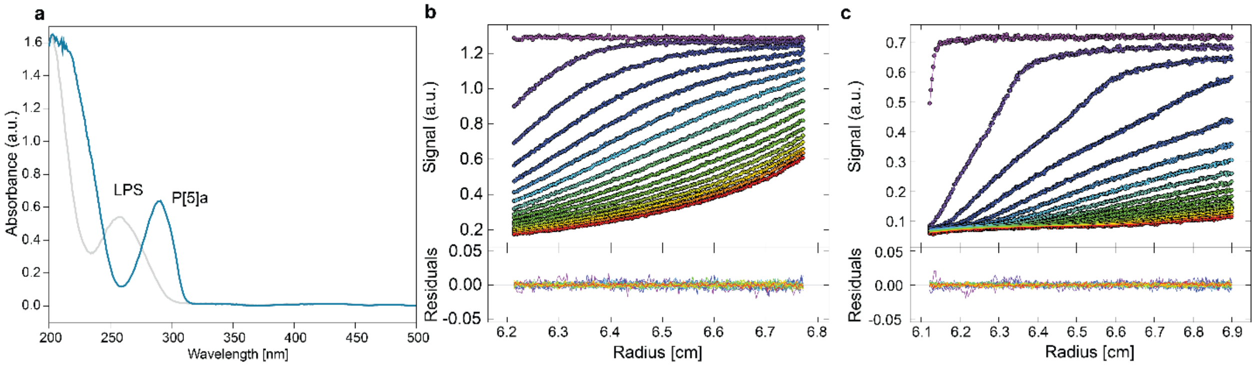

**Supplementary Figure 9.** Absorbance and sedimentation spectra of P[5]a and LPS. **a**) Absorbance spectra of P[5]a and LPS show wavelengths that allow monitoring of individual compounds in the mixture (260 nm LPS, 305 nm P[5]a). At 260 nm, the absorption peak of LPS, the absorbance of P[5]a is negligible. Conversely, at 305 nm, the absorbance of LPS is negligible. **b**) Absorbance sedimentation boundaries of 0.5 g $L^{-1}$ LPS in 0.5x PBS buffer at 60.000 rpm (circles) and best-fit solution by c(s) model (lines). **c**) Absorbance sedimentation boundaries of 0.5 g $L^{-1}$ LPS in 0.5x PBS buffer at 60.000 rpm (circles) and best-fit solution by c(s) model (lines). Color gradient represents the time from beginning of the experiment (violet) to the end (red).

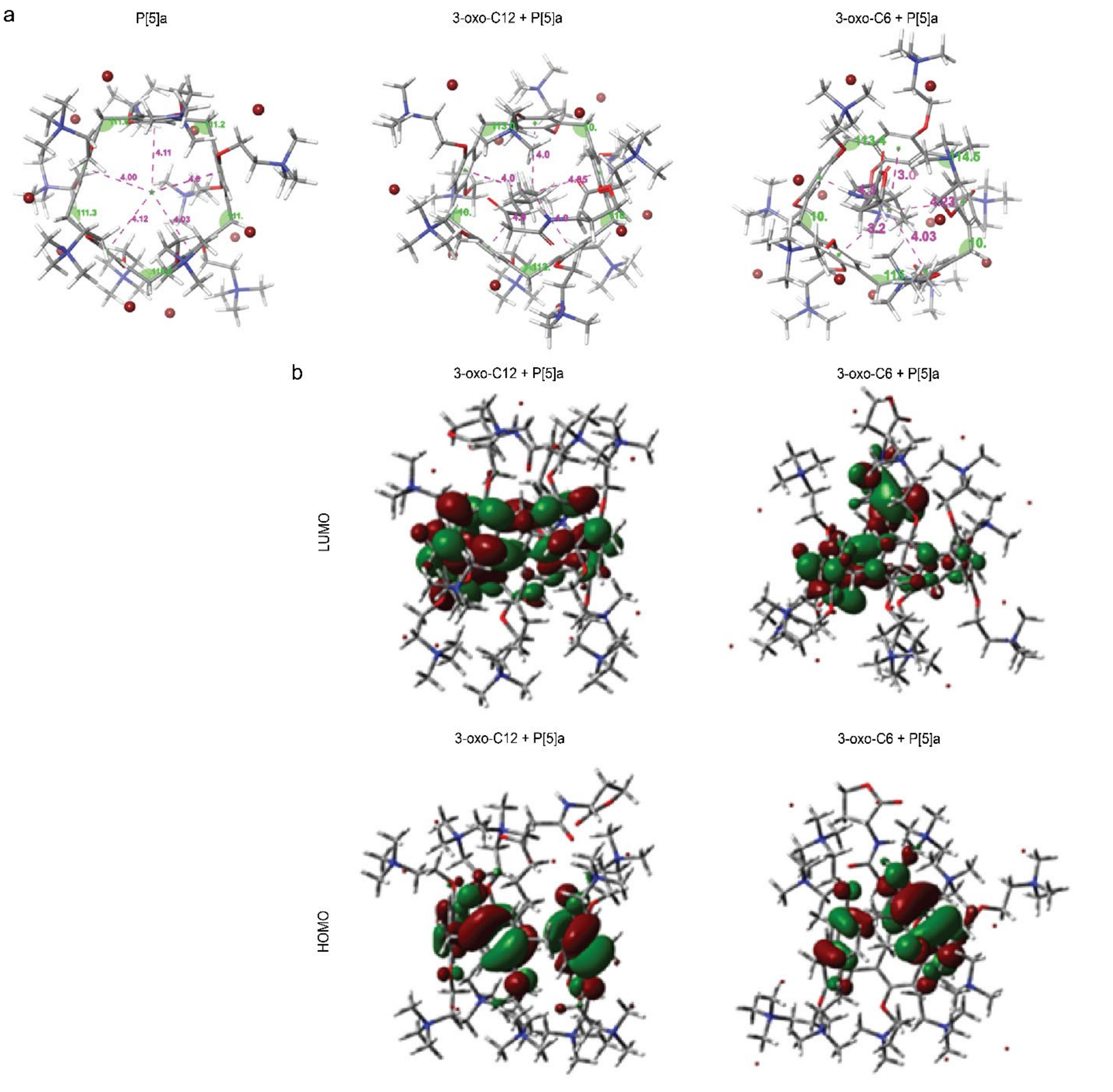

**Supplementary Figure 10. a**) The predicted distances between the aromatic rings and the centre of the cavity, and the angle between adjacent aromatic rings in the isolated host structure versus the host within complex geometries of 3-oxo-C12 and P[5]a, and 3-oxo-C6 and P[5]a. **b**) 3D representations of the HOMO and LUMO orbitals within the M062X/6-311G**calculated complex structures of 3-oxo-C12 and P[5]a, and 3-oxo-C6 and P[5]a.

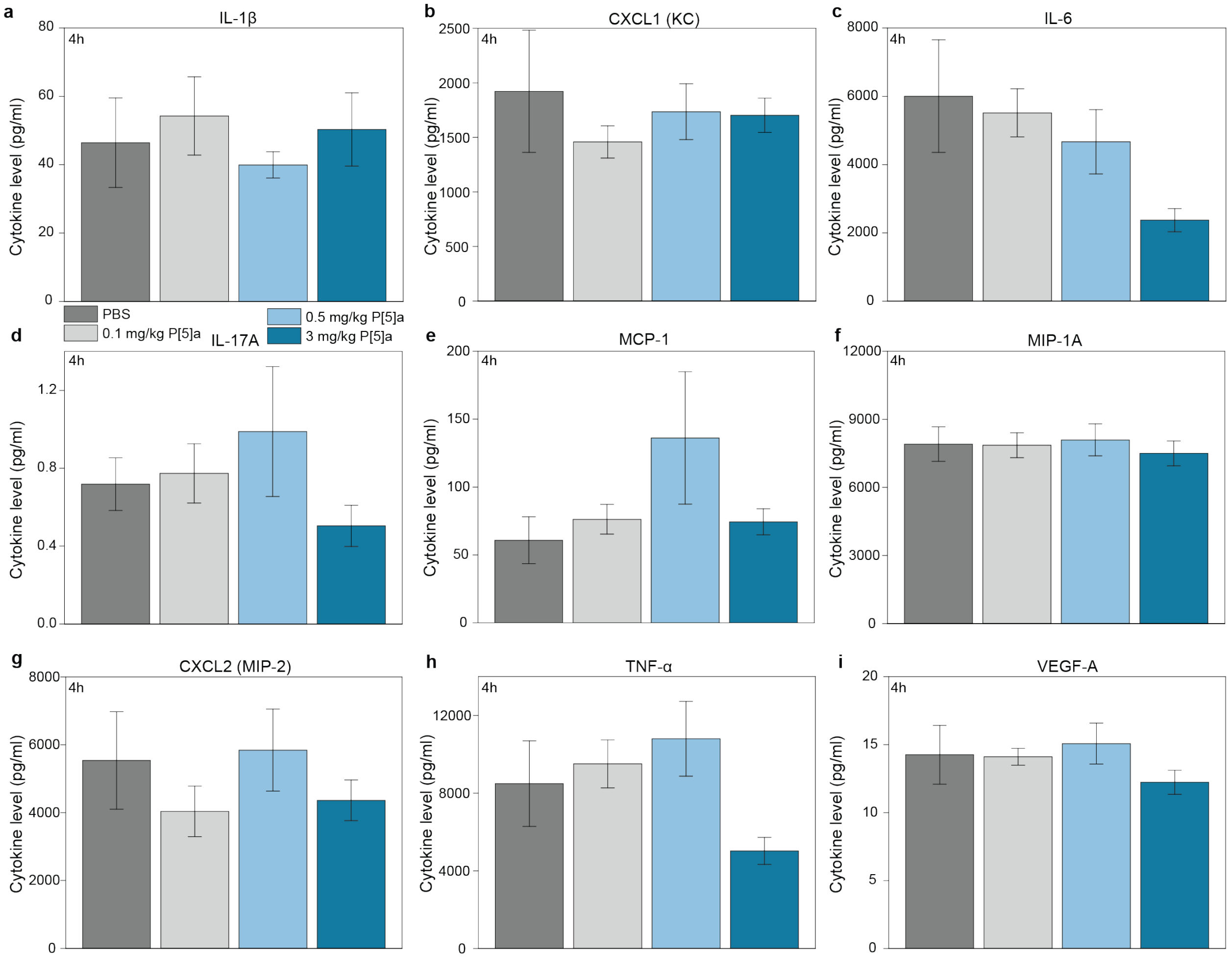

**Supplementary Figure 11.** Effects of P[5]a, dosed i.t., on LPS-stimulated (**a**) IL-1β, (**b**) CXCL1 (KC), (**c**) IL-6, (**d**) IL-17A, (**e**) MCP-1, (**f**) MIP-1A, (**g**) CXCL2 (MIP-2), (**h**) TNF-α and (**i**) VEGF-A counts in BALF at 4h. One animal (3 mg/kg, 4h) died immediately after i.t. treatments. Statistical analysis was performed by One Way ANOVA followed by Dunnett's test for multiple comparison, and Grubb's test was performed to exclude outliers. Mean (± SEM) (n=10).

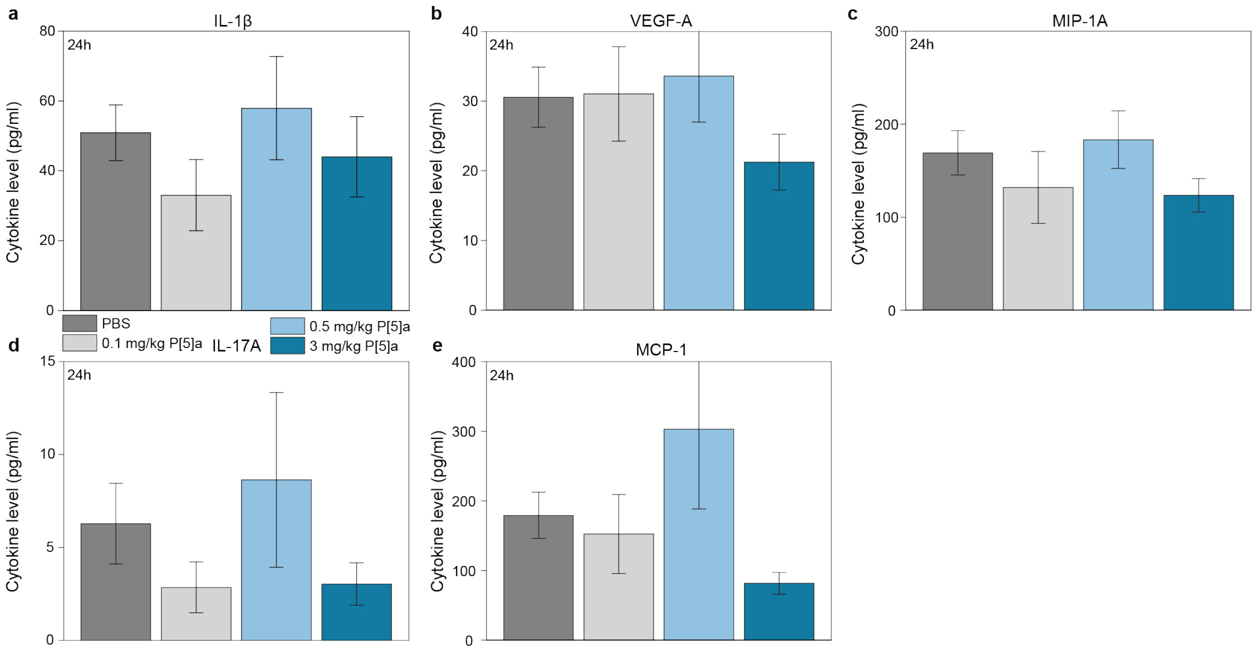

**Supplementary Figure 12.** Effects of P[5]a, dosed i.t., on LPS-stimulated (**a**) IL-1β, (**b**) VEGF-A, (**c**) MIP-1A, (**d**) IL-17A and (**e**) MCP-1 counts in BALF at 24h. One animal (3 mg/kg, 4h) died immediately after i.t. treatments. Statistical analysis was performed by One Way ANOVA followed by Dunnett's test for multiple comparison. Mean (± SEM) (n=10).

P[5]a synthesis

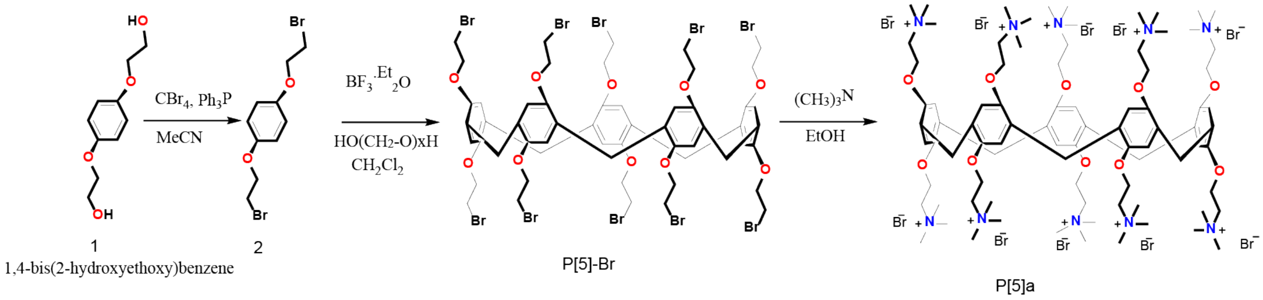

**Supplementary Figure 13.** 3-step synthesis of Pillar[5]arene bromide.

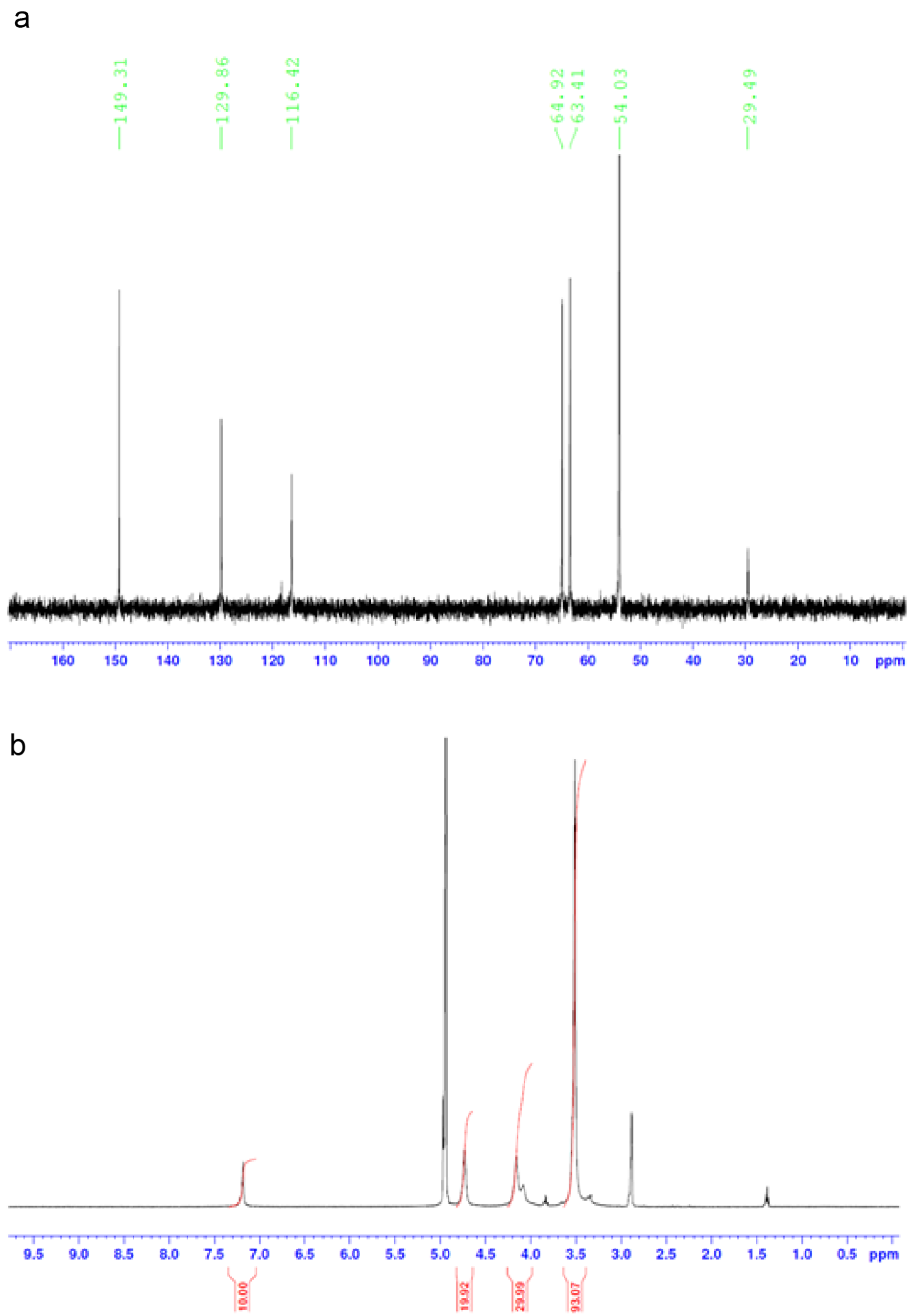

**Supplementary Figure 14. a**) Pillar[5]arene-13C NMR in $D_2O$. **b**) Pillar[5]arene-1H NMR in $D_2O$-DMSO.

a

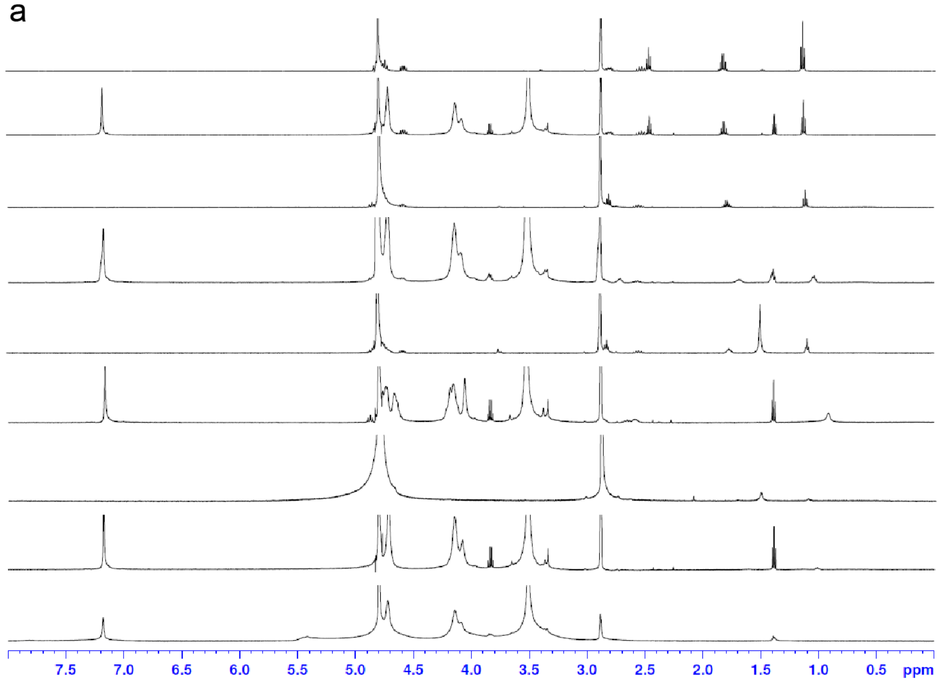

b

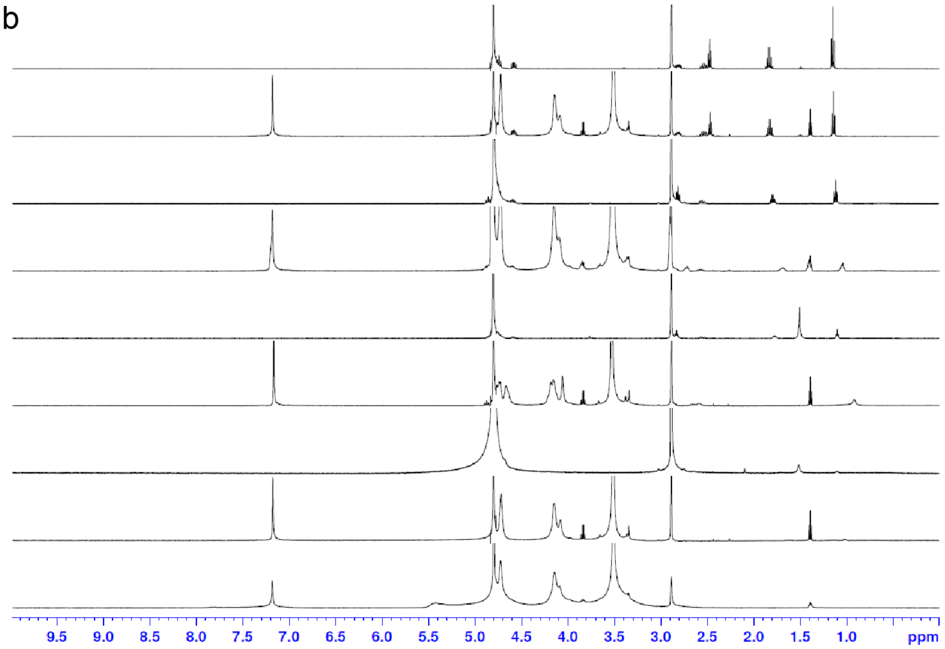

**Supplementary Figure 15. a**) Pillar[5]arene-1H NMR in $D_2O$ 8.0-ppm. **b**) Pillar[5]arene-1H NMR in $D_2O$-10.0 ppm. **b**) COSY spectrum of P[5]a in DMSO-d6.

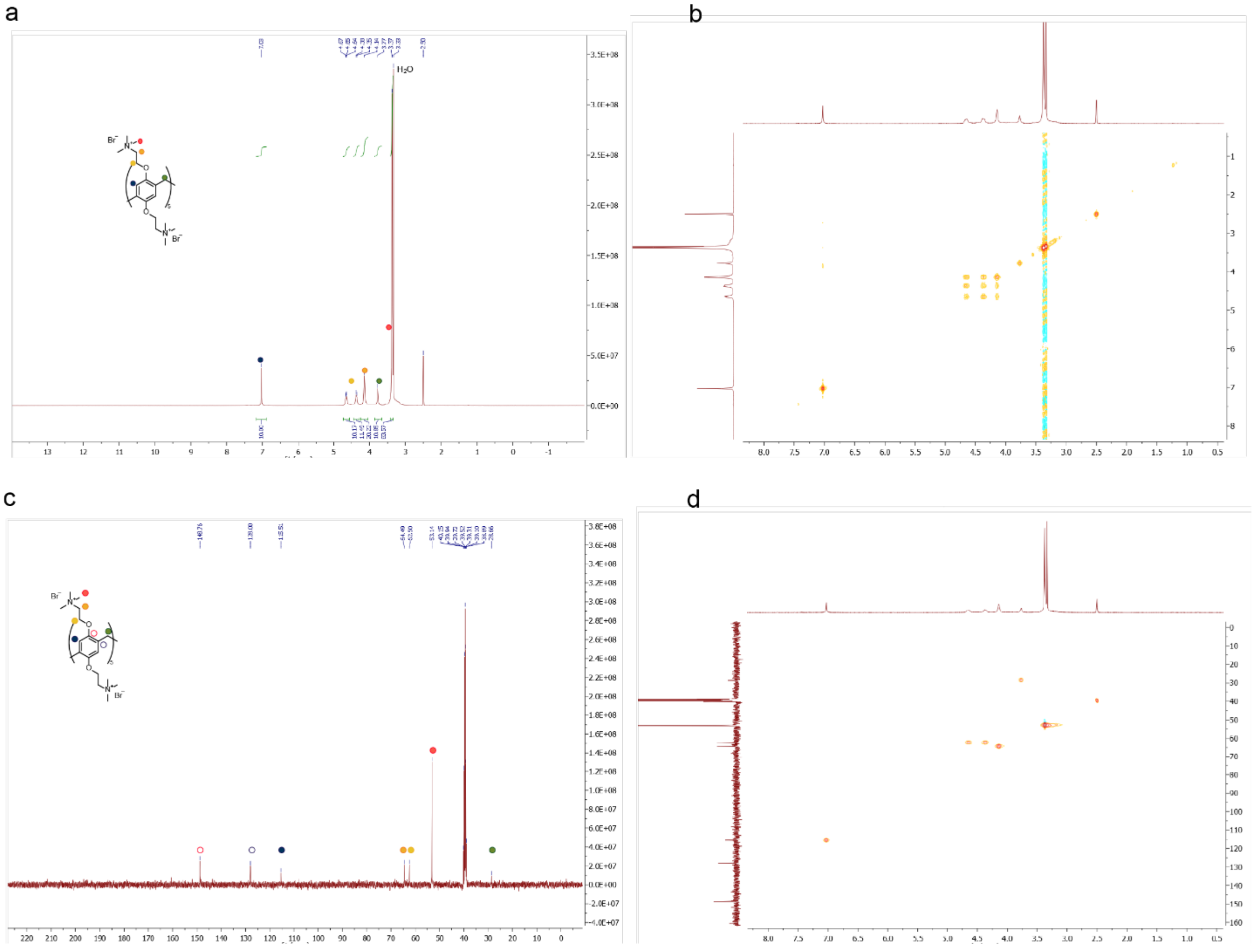

**Supplementary Figure 16.** **a**)[1]H-NMR spectrum of dried overnight P[5]a in DMSO-d6 (400 MHz), including chemical structure, assigned peaks, and residual solvent signals. **b**) COSY spectrum of dried overnight P[5]a in DMSO-d6. **c**) $^{13}$C-NMR spectrum of dried overnight P[5]a in DMSO-d6 (125 MHz), including chemical structure, assigned peaks, and residual solvent signals. **d**) HSQC spectrum of dried overnight P[5]a in DMSO-d6.

### Supplementary Methods

#### RNA-sequencing

RNA was extracted from the A549 cells incubated under different conditions using an RNeasy Mini Kit (Qiagen) following manufacturer's guidelines. Cells were homogenized with QIAshredder columns (Qiagen). After extraction, RNA was quantified using a Qubit 2 fluorometer (Thermo Fisher Scientific) with the Qubit RNA HS Assay Kit (Thermo Fisher Scientific), and stored at -70°C. The RNA sequencing method was designed based on the Drop-seq protocol described in[9].

Briefly, 10 ng of RNA was mixed with Indexing Oligonucleotides (Integrated DNA Technologies). After 5 minutes of incubation at ambient temperature, RNA was combined with RT mix containing 1× Maxima RT buffer, 1 mM dNTPs, 10 U $\mu L^{-1}$ Maxima H- RTase (all Thermo Fisher Scientific), 1 U $\mu L^{-1}$ RNase inhibitor (Lucigen) and 2.5 µM Template Switch Oligo (Integrated DNA Technologies). Samples were incubated in a T100 thermal cycler (BioRad) for 30 minutes at 22°C, then 90 minutes at 52°C. The constructed cDNA was amplified by PCR in a volume of 15 µL using 5 µL of RT mix as the template, 1× HiFi HotStart Readymix (Kapa Biosystems) and 0.8 µM SMART PCR primer. The thermocycling profile was as follows: 95°C for 3 min; four cycles of 98°C for 20 sec, 65°C for 45 sec, 72°C for 3 min; 13 cycles of 98°C for 20 sec, 67°C for 20 sec, 72°C for 3 min; and final extension step of 5 min at 72°C. The PCR products were pooled together in sets of nine samples containing different Indexing Oligos, purified with 0.6× Agencourt AMPure XP Beads (Beckman Coulter) according to manufacturer's instructions, and eluted in 10 µL of molecular grade water. The 3'-end cDNA fragments for sequencing were prepared using the Nextera XT (Illumina) tagmentation reaction with 600 pg of each PCR product serving as an input. The reaction was performed according to manufacturer's instructions, with the exception of the P5 SMART primer that was used instead of the S5xx Nextera primer. Each set of 12 samples that was pooled after the PCR reaction was tagmented with a different Nextera N7xx index. The samples were then PCR amplified as follows: 95°C for 30 sec; 11 cycles of 95°C for 10 sec, 55°C for 30 sec, 72°C for 30 sec; and a final extension step of 5 min at 72°C. Samples were purified twice using 0.6× and 1.0× Agencourt AMPure Beads (Beckman Coulter), and eluted in 10 µl of molecular grade water.

The overnight culture of *P. aeruginosa* PAO1 was diluted 1:100 in fresh LB medium and incubated aerobically for 3 h at 37°C. The culture was then diluted again 1:100 and grown in a total volume of 200 µL with or without the addition of 2.5 mM P[5]a in a 96-well plate. Bacterial cultures were incubated for 24 h with constant shaking. Two 200 µL cultures were pooled together and the total RNA was isolated from three samples using an RNeasy Mini Kit (Qiagen) following manufacturer's guidelines. The rRNA was removed using a Ribo-ZeroTM rRNA Removal kit for Bacteria (Epicentre). The first strand cDNA was constructed using the SuperScript II Reverse Transcriptase (Thermo Fisher Scientific), while the second strand was constructed using random hexamers (Thermo Fisher Scientific) and HiFi HotStart Readymix (Kapa Biosystems). The sequencing library was prepared using the Nextera XT (Illumina) tagmentation reaction with 1 ng of RNA serving as an input, following manufacturer's instructions. Samples were purified twice using 0.6× and 0.9× Agencourt AMPure Beads (Beckman Coulter), and eluted in 10 µL of ultra-pure water.

The concentration of both libraries was measured using a Qubit 2 fluorometer (Thermo Fisher Scientific) and a Qubit DNA HS Assay Kit (Thermo Fisher Scientific). The quality of the sequencing libraries was assessed using the LabChip GXII Touch HT electrophoresis system (PerkinElmer), with the DNA High Sensitivity Assay (PerkinElmer) and DNA 5K / RNA / Charge Variant Assay LabChip (Perkin Elmer). Samples were stored at -70°C. The libraries were sequenced on an Illumina NextSeq 500, with the addition of the custom primer, producing read 1 of 20 bp and read 2 (paired end) of 55 bp. Sequencing was performed at the Functional Genomics Unit of the University of Helsinki, Finland. RNA sequencing data has been deposited to NCBI Gene Expression Omnibus (Accession ID: GEO Submission GSE182853).

**Read alignment and RNA-seq data analysis**

**A549 cells.** Raw sequence data was filtered to remove reads shorter than 20 bp. The original pipeline suggested for processing drop-seq data was used. Briefly, reads were additionally filtered to remove polyA tails 6 bp or longer, then aligned to the human (GRCh38) genome using STAR aligner[17] with default settings. Uniquely mapped reads were grouped according to the 1-9 barcode, and gene transcripts were counted by their Unique Molecular Identifiers (UMIs) to reduce bias emerging from PCR amplification. Digital expression matrices (DGE) reported the number of transcripts per gene in a given sample (according to the distinct UMI sequences counted). Differentially expressed genes were identified using DESeq2[18] with the cut-off for the adjusted p-value set to 0.001. A heatmap of gene expression levels was created using Heatmapper (http://www1.heatmapper.ca/). Venn diagrams were created to evaluate the distribution of differentially expressed genes between specified groups using Venny (http://bioinfogp.cnb.csic.es/tools/venny/).

***P. aeruginosa***. The bacterial sequencing reads were filtered for quality and aligned against the *P. aeruginosa* PAO1 genome (accession number NC_002516.2) using the BowTie2 read aligner[19]. DESeq2[20] was then used to obtain the list of differentially expressed genes. Genes were considered differentially expressed if the log2 fold change was > ±2 and the adjusted *P* value was < 0.001. RNA sequence data has been deposited to Gene Expression Omnibus (Accession ID: GEO Submission GSE182847).

**Supplementary methods Computational Discussion**

Density Functional Theory (DFT) was used to explore the interactions of 3-oxo-C12 and 3-oxo-C6 with the P[5]a receptor. The M062X/6-311G** level of theory, appropriate for supramolecular interactions due to its handling of weak non-covalent interactions, was employed[10,11,21]. This choice was validated as dispersion-uncorrected functionals (i.e. B3LYP) failed to provide meaningful data. The HSLs show good surface-size complementarity with P[5]a; as the total binding surface area is generally proportional to the strength of a binding interaction, this is consistent with our expectations and the long hydrophobic chains all reside in the internal cavity which adjusts to the size of the encapsulated guest. The lactone and the neighbouring hydrophilic functionalities interact with the rim-functionalities of the P[5]a molecule. The 3-Oxo-C12 with P[5]a complexes show far higher affinity than any other pair examined in the study.

**Total binding-surface areas**

Electron density isosurface values.

The electron density best approximates the size and shape of the molecule, thus larger molecules with more electrons feature larger isosurface values than smaller molecules with fewer electrons. As shown by the calculated density isosurfaces (Supplementary Fig. 7), the pillararene's cylinder closes around 3-oxo-C12 with P[5]a with Electron density from total SCF density (npts = 71,68,71; res (A) = 0.344690, 0.344690, 0.344690) plus 3-oxo-C12 fragment [(Electron density from Total SCF Density (npts = 71,68,71; res (A) = 0.344690, 0.344690, 0.344690). A far smaller potential binding surface for both systems is observed for 3-oxo-C6 with P[5]a [(Electron density from Total SCF Density (npts = 74,72,64; res (A) = 0.387932, 0.387932, 0.387932) plus 3-oxo-C6 Fragment [(Electron density from Total SCF Density (npts = 74,72,64; res (A) = 0.387932, 0.387932, 0.387932).

**Hydrophobic/philic surface areas**

Hydrophilic regions are shown by contouring the "hydrophilic grid" at a negative isosurface value of -6 kcal/mol (blue surfaces) while the hydrophobic regions are shown by contouring the associated grid at a suitably negative threshold of −0.5 kcal/mol (orange surfaces). These surfaces include both the interactions between the two components, and their interaction with solvent (especially in the case of the host). Note that 3-oxo-C12 shows an extensive series of hydrophobic interaction within the cavity, and hydrophilic interactions at the mouth of the cavity due to the interaction between the lactone and the nearby functional groups with the ammonium rim groups of the pillararene. 3-oxo-C6, on the other hand, has the entire HSL inserted into the tunnel removing these beneficial interactions meaning the guest has very few interactions with the host (Supplementary Fig. 8). This is accompanied by a disruption of the external interactions for the host: the surface is far more disrupted meaning that there is less possible interaction between host and solvent than in either of the other two systems. These distortions and weak interactions help explain the comparably low affinity of 3-oxo-C6 for the host compared to the 3-oxo-C12.

**The structural parameters**

Binding is facilitated when the incorporation of the guest does not induce a significant distortion in the preferred conformation of the host. This can be quantified by looking at the relative difference in shape between the complexes and the isolated systems. As will be discussed below in 6.3, electronic distortions follow a similar principle.

For these systems, we can quantify the disruption most clearly by calculating the centroid distances in the cavity (the distances between the aromatic rings and the centre of the cavity), and torsional strain induced on the pillararene (the angle between adjacent aromatic rings; Supplementary Fig. 8d). In the isolated cavity, prepared using our computational model, the distances tend to be 4.0-4.1 Å, and the bond angle 110º. This is identical to the values observed in the crystal structure.

Along with the non-specific hydrophobic interactions, various strong, defined intramolecular interactions are responsible for the high affinity. These can be quantified as observed by the plotted topological molecular graph showing bond critical points and bond paths using quantum theory of atoms in molecules (QTAIM). The analysis is provided at Supplementary Fig. 8c. An extended table of the electronic charge density ($\rho_{BCP}$) and its Laplacian ($\nabla^2\rho$) at the bond critical point (BCP) representing both the intra-host and host-guest non-covalent interactions is provided below as Supplementary table 4 for 3-oxo-C12. The interactions (e.g., H-bonding network and noncovalent interactions) responsible for guest binding are larger in number and all stronger in the 3-oxo-C12 with P[5]a than 3-oxo-C6 with P[5]a, as confirmed by the predicted larger interaction energy since the larger portion of the hydrophobic terminal part of the guest is positioned inside the binding cavity and involved in the formation of the attractive interactions. Notably, the relative energetic differences between 3-oxo-C12 with P[5]a and 3-oxo-C6 with P[5]a became larger upon using of the hybrid dispersion corrected DFT based functional (i.e., M062X) versus the dispersion-uncorrected functional (i.e., B3LYP) which further highlights the importance of these interactions in obtaining high selectivity and binding affinity. Multiple stabilizing $CH_{alkyl}\cdots\pi_{Ar}$ interactions inside the bowl-shaped cavity of P[5]a are highlighted, and their attractive contribution is confirmed by the presence of the (3, −1) critical points (BCP) between the bond path linking the guest's hydrogen atoms of long chain with aromatic carbon atoms of the host, with the electron density ($\rho_{BCP}$) in the range of around 0.0040−0.0080 au for 3-oxo-C12 with P[5]a complexes. The Laplacian of the charge density $\nabla^2\rho$ ranging from 0.013 to 0.020 au for the related interactions, are all positive showing the closed-shell (*i.e.* ionic bonds, H-bonds and van der Waals interactions) character of these interactions. Additionally, the intermolecular $CH_{alkyl}\cdots O_{Ar}$ non-classical type H-bonding interactions with BCP ($\rho_{BCP}$ = 0.003 to 0.0098 au) and ($\nabla^2\rho$ = 0.010 to 0.033) between the phenolic oxygen atoms of P[5]a leads to even larger stability of 3-oxo-C12 with P[5]a complexes compared with 3-oxo-C6 with P[5]a. The energetically unfavorable 3-oxo-C6 with P[5]a complex did not show stabilization interactions with the same strength.

Supplementary Table 4. The predicted QTAIM $\rho_{BCP}$ and $\nabla^2\rho$ representing the host-guest non-covalent interactions were obtained from the M062X/6-311G**electron density for the favored P[5]a with 3-oxo-C12 complexes.

| | $\rho_{BCP}$ ($e/au^3$) | $^2\rho$(r) ($e/au^3$) |
|---|---|---|
| $CH_{Alkyl}...\pi_{Ar}$ | 0.007 | 0.017 |
| | 0.007 | 0.017 |
| | 0.005 | 0.013 |
| | 0.005 | 0.015 |
| | 0.006 | 0.016 |
| | 0.006 | 0.016 |
| | 0.004 | 0.010 |
| | 0.004 | 0.010 |
| | 0.006 | 0.016 |
| | 0.004 | 0.010 |
| | - | - |
| $CH_{Alkyl}...O_{Ar}$ | | |
| | 0.006 | 0.016 |
| | 0.008 | 0.021 |
| | 0.011 | 0.034 |
| | 0.003 | 0.011 |
| | 0.004 | 0.013 |
| | - | - |
| | - | - |
| | - | - |

**The electronic parameters**

Absolute, and shifts in, the electrostatic charge distributions

To evaluate electrostatic interactions, we employed differential electrostatic potential (dESP) maps overlaid on electron isodensity contours to visualize both absolute and shift in the electrostatic charge distribution upon complexation (Supplementary Fig. 7). In isolated P[5]a, the positive charge potential (blue) is located mostly on the rim ammonium functionalities, and the negative charge potential (red) is distributed on the outer and inner face of the pillararene cavity as well as on the counterions; however the degree of charge separation is minimal. The isolated 3-oxo-C12 and 3-oxo-C6, are largely neutral (green) with only slight charge separation on the headgroup.

Upon complexation the system changes significantly, inducing shifts in electron density to maximize complex stability. The differences between the systems can be readily identified. Association with 3-oxo-C12 adds electron density to the ammonium functionalities, making the terminals of the cavity less polarized than in the isolated system. In contrast, 3-oxo-C6 dramatically distorts the electronics of the system as demonstrated by the less organized, more numerous, and more intense blue and red surfaces: the host is not stabilized by the presence of the guest. This is best illustrated by the difference maps. In these, 3-oxo-C12 induces a relative negative shift in the ammonium groups (as electron density from the guest is transferred to these charged elements). For 3-oxo-C6 the surface is far more compromised with multiple different areas shifting density demonstrating the large structural distortions to the preferred electronic distribution that occur to accommodate 3-oxo-C6; this relatively greater distortion is yet another reason for the lower affinity of this system.

Molecular Orbital Theory analysis:
To determine the change transfer and the electronic contributions of complexation process, the frontier molecular orbitals were calculated for all complexes. We find that the energy differences between the complexes are minor in all cases, the frontier molecular orbital component of the binding energy is favourable as the HOMO-LUMO gap decreases upon complexation, but the differences between the systems are minimal for both host ($\Delta\Delta E_{HOMO\text{–}LUMO(Host)}$= -0.09, and -0.17 eV for P[5]a upon complexation with 3-oxo-C12 and 3-oxo-C6 respectively) and guest ($\Delta\Delta E_{HOMO\text{–}LUMO(Guest)}$= -0.40, -0.34 eV for 3-oxo-C12, and 3-oxo-C6 respectively upon complexation to P[5]a). Furthermore, the resultant orbital density localization of the complexes are similar: charge transfer evaluated form FMO are not a discriminating factor but do help drive complexation in all cases (Supplementary Fig. 10).
